# Sacsin cotranslational degradation causes autosomal recessive spastic ataxia of Charlevoix-Saguenay

**DOI:** 10.1101/2021.03.16.435646

**Authors:** Fabiana Longo, Daniele De Ritis, Annarita Miluzio, Davide Fraticelli, Jonathan Baets, Marina Scarlato, Filippo M. Santorelli, Stefano Biffo, Francesca Maltecca

**Affiliations:** Mitochondrial Dysfunctions in Neurodegeneration Unit, Ospedale San Raffaele, Milan, Italy; Istituto Nazionale di Genetica Molecolare, INGM, “Romeo ed Enrica Invernizzi”, Milan, Italy; Translational Neurosciences, Faculty of Medicine and Health Sciences, UAntwerpen, Antwerp, Belgium; Laboratory of Neuromuscular Pathology, Institute Born-Bunge, University of Antwerp, Antwerpen, Belgium; Neuromuscular Reference Centre, Department of Neurology, Antwerp University Hospital, Antwerpen, Belgium; Department of Neurology, Ospedale San Raffaele, Milan, Italy; Molecular Medicine, IRCCS Fondazione Stella Maris, Pisa, Italy; Department of Biosciences, University of Milan, Milan, Italy; Università Vita-Salute San Raffaele, Milan, Italy

**Keywords:** autosomal recessive spastic ataxia of Charlevoix-Saguenay, genotype-phenotype correlation, molecular mechanisms of pathogenesis

## Abstract

Autosomal recessive spastic ataxia of Charlevoix-Saguenay is caused by more than 200 different mutations in the *SACS* gene encoding sacsin, a huge multimodular protein of unknown function. ARSACS phenotypic spectrum is highly variable. Previous studies correlated the nature and position of *SACS* mutations with age of onset or disease severity, though the effects on protein stability were not considered.

In this study, we explain mechanistically the lack of genotype-phenotype correlation in ARSACS, with important consequences for disease diagnosis and treatment.

We found that sacsin is almost absent in ARSACS fibroblasts, regardless of the nature of the mutation. We did not detect sacsin in patients with truncating mutations, while we found it strikingly reduced or absent also in compound heterozygotes carrying diverse missense mutations. We excluded *SACS* mRNA decay, defective translation, or faster post-translational degradation as causes of protein reduction. Conversely, we demonstrated that nascent mutant sacsin protein undergoes preemptive cotranslational degradation, emerging as a novel cause of a human disease. Based on these findings, sacsin levels should be included in the diagnostic algorithm for ARSACS.

## Introduction

ARSACS is an early onset neurological disease first described in Québec (Canada), due to a founder effect (Bouchard et al., 1978), and is one of the most frequent recessive ataxias after the Friedreich’s ataxia. Clinically, ARSACS is characterized by progressive cerebellar ataxia, spasticity and sensorimotor peripheral neuropathy. Most patients present with the above triad of symptoms, typically with early onset cerebellar ataxia followed by spasticity and later by neuropathy. However, the clinical spectrum is highly variable among patients, with an increasing number of diagnosed subjects with disease onset in early adult-years, or with a clinical presentation including only one or two of the three typical symptoms. Also, a minority of patients present intellectual disability, epileptic seizures, urinary dysfunction and hearing loss (Baets et al., 2010; Synofzik et al., 2013; Vermeer et al., 1993 updated 2020; Xiromerisiou et al., 2020).

ARSACS is caused by mutations in the *SACS* gene, which is one of the largest of our genome with a gigantic exon 10 of 12.8 kb (Engert et al., 2000) and nine smaller upstream exons. *SACS* gene encodes the 520 kDa multimodular protein sacsin, highly expressed in the central nervous system and in particular in the cerebellum (Lariviere et al., 2019).

From the N-terminus to the C-terminus, the amino acid sequence of sacsin contains: an ubiquitin-like (UbL) domain that binds to the proteasome (Parfitt et al., 2009), three sacsin-repeating regions (SRR) having high homology with Hsp90 (Anderson et al., 2010), a Xeroderma Pigmentosus group C protein Binding (XPCB) domain (Greer et al., 2010), a DnaJ domain that binds Hsc70 (Parfitt et al., 2009) and a Higher Eukaryotes and Prokaryotes Nucleotide-binding (HEPN) domain (Kozlov et al., 2011). Despite the nature of these motifs suggests that sacsin may operate in protein quality control, the cellular function of this protein and the pathophysiological consequences of its dysfunction remain largely unknown.

More than 200 mutations have been described worldwide to date, almost equally divided in homozygous or compound heterozygous (Xiromerisiou et al., 2020). The majority are missense, followed by small deletions, frameshift and nonsense and spread over the whole gene, as expected for a recessive disease. No clear-cut genotype-phenotype correlations have been identified in ARSACS so far (Baets et al., 2010; Bouhlal et al., 2011; Synofzik et al., 2013; Vermeer et al., 1993 updated 2020). Patients with macrodeletions that lead to loss of more than 3000 amino acids have similar phenotype to those that harbor single base insertion/deletions (indels) in the C-terminal end of the protein or that have missense substitutions (Bouhlal et al., 2011). Moreover, there are also reports of inter- or intrafamilial variability in patients with the same mutations (Bouhlal et al., 2011; Gagnon et al., 2018; Krygier et al., 2017).

Some studies have been published trying to correlate the pathogenicity and/or the position of the *SACS* mutations with the age of onset or the clinical severity of the disease. By analyzing genetic and clinical data from 70 ARSACS patients, Romano *et al.* suggested that subjects carrying missense mutations in homozygosity or heterozygosity with a null allele exhibit significantly milder phenotypes (evaluated by an arbitrary clinical score) than those carrying a truncating mutation on each allele. Moreover, they propose that mutations in the N-terminal part of the SRR domains impact more severely on ARSACS phenotype compared to those in the C-terminal part (Romano et al., 2013). In a more recent work, considering all the published ARSACS mutations up to 2019, Xiromerisiou *et al.* identified a correlation between the pathogenicity score of the mutations (calculated by computational algorithms), the phenotype severity and the age of onset (Xiromerisiou et al., 2020). However, these *in silico* studies have major limitations, since they did not consider the consequences of the various *SACS* mutations on sacsin protein levels.

In this work, by employing both commercial and newly developed anti-sacsin antibodies, we demonstrated that sacsin is almost absent in a large panel of ARSACS fibroblasts, regardless of the nature and position of the mutation along the gene. We indeed did not detect sacsin (either full-length protein or truncated products) in patients with null mutations. Unexpectedly, we found it strikingly reduced (or completely absent) in compound heterozygotes carrying missense mutations. We excluded faster post-translational degradation of sacsin with missense mutations, as inhibition of cellular degradative pathways, as well as blockade of different classes of proteases, never rescued sacsin levels in ARSACS fibroblasts. Also, we detected comparable or even higher sacsin mRNA levels in patients with missense mutations compared to controls, excluding mRNA degradation. Polysome profiling revealed no evident defects in sacsin translation in ARSACS patients. Our data demonstrate that sacsin upon missense mutations undergoes cotranslational ubiquitination and degradation, and this event prevents the complete full-length sacsin production. To our knowledge, this is the first report in which this mechanism, still poorly characterized, has been formally demonstrated *in vivo* for a mammalian protein with pathogenetic mutations.

Overall, our data indicate that the phenotypic variability in ARSACS is not due to the different levels of residual sacsin or positional effect of the mutations. We provide evidence of a new molecular explanation of the lack of genotype-phenotype correlation in ARSACS patients, defining the loss of sacsin protein as a unifying mechanism shared by the different *SACS* mutations.

## Results

### Full-length sacsin protein is dramatically reduced in ARSACS patient fibroblasts regardless of the nature of the mutation

To study the impact of *SACS* mutations on sacsin levels, we derived skin fibroblasts from eleven compound heterozygous ARSACS patients, for a total of nineteen different mutations analyzed (**Fig. 1**). Detailed clinical features of patients are reported in **Table 1**. To have a broad picture of ARSACS genetics, we selected patients carrying mutations of different nature and localized in different domains of sacsin: compound heterozygous for a missense on one allele and a truncating frameshift on the other allele (PN1, PN2, PN7), compound heterozygous for different truncating mutations (PN3, PN6a, PN6b), compound heterozygous for a missense mutation and a big deletion (PN4), compound heterozygous for truncating mutations and a big deletion (PN8a, PN8b), compound heterozygous for missense mutations only (PN5, PN9) (**Fig. 1**). PN6a and PN6b, and PN8a and PN8b are siblings, and despite carriers of the same mutation, they present phenotypic variability (**Table 1**).

**Table 1.**
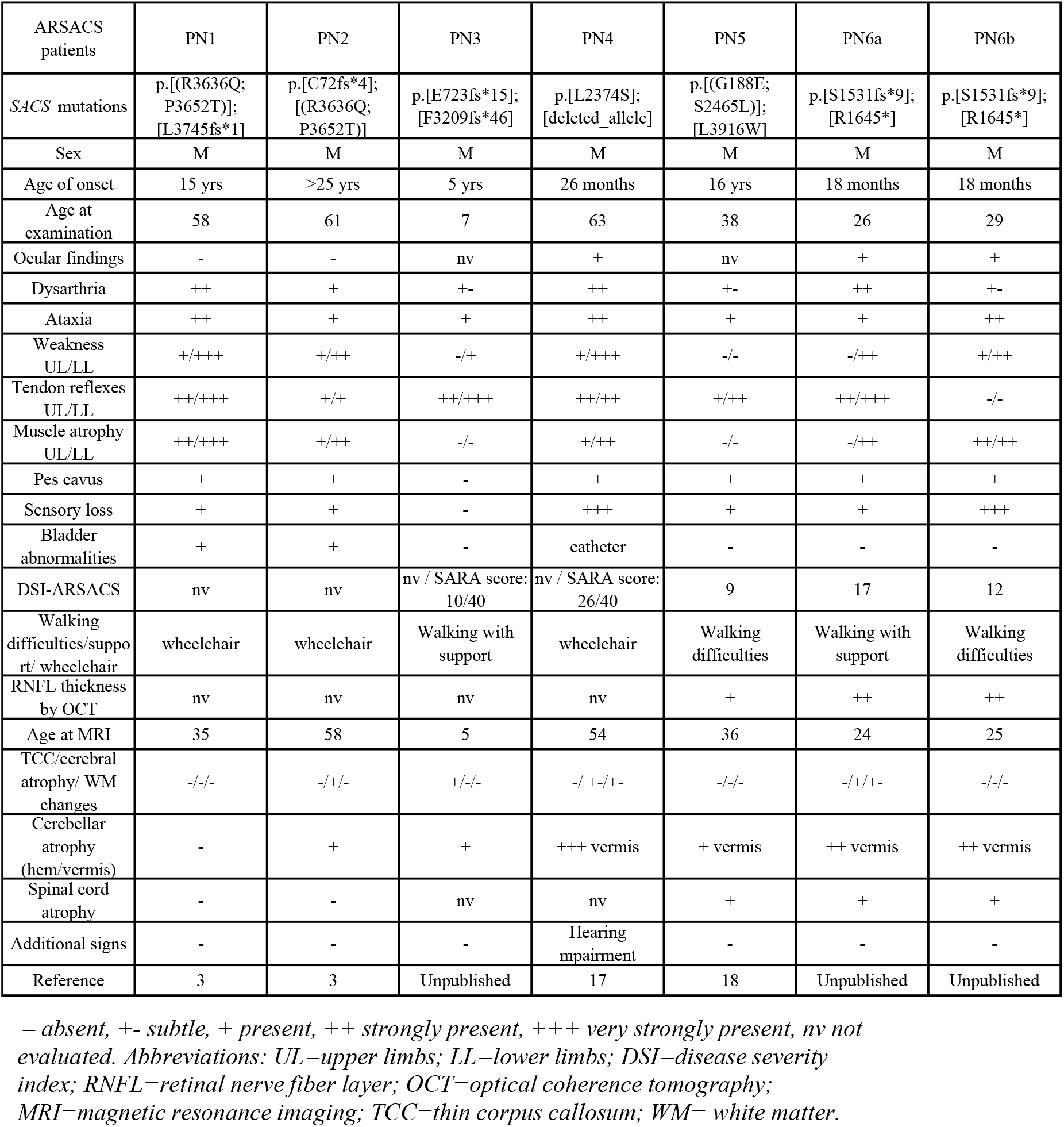
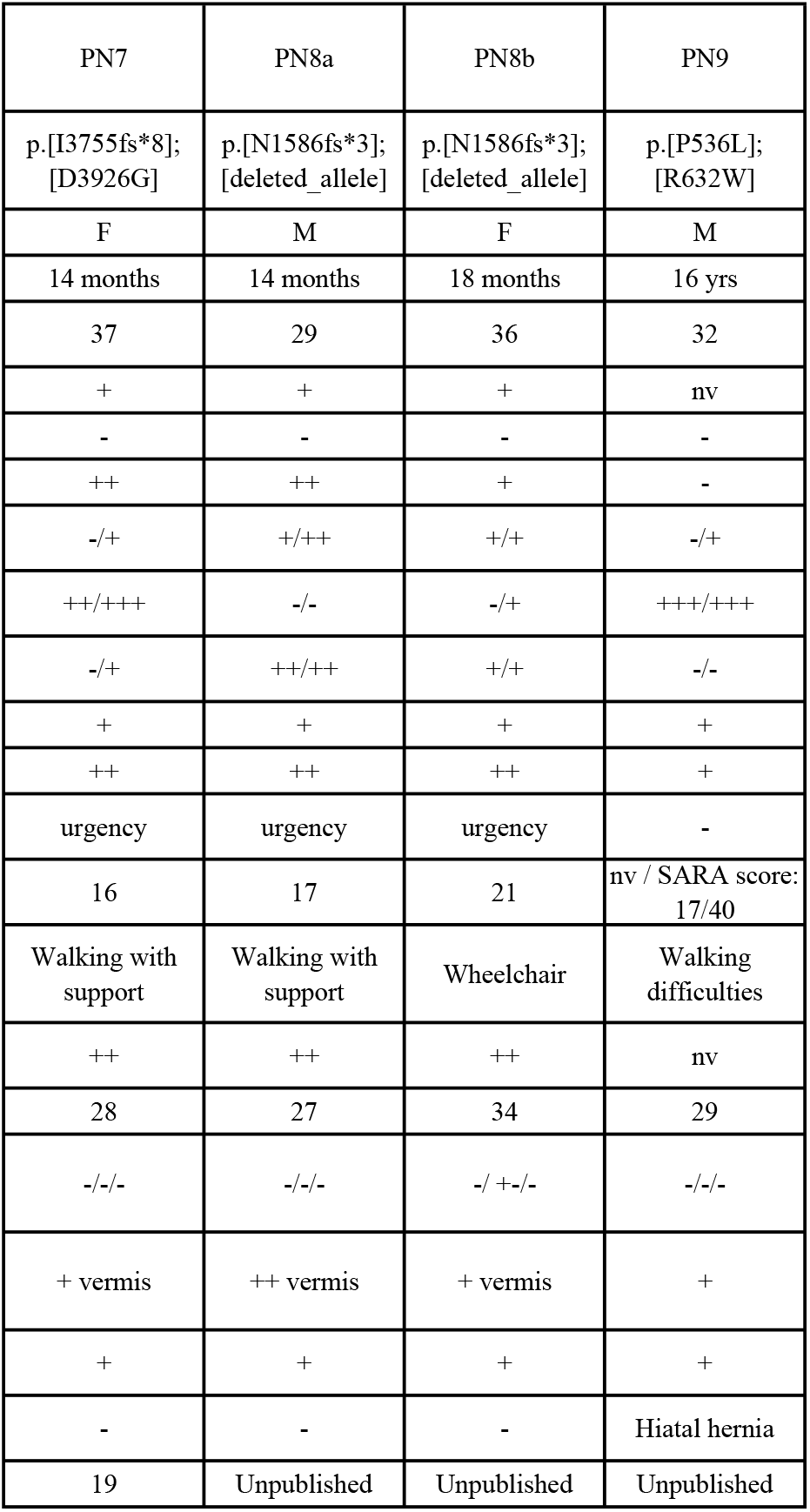
Clinical features of ARSACS patients reported in this study.

**Figure 1.**
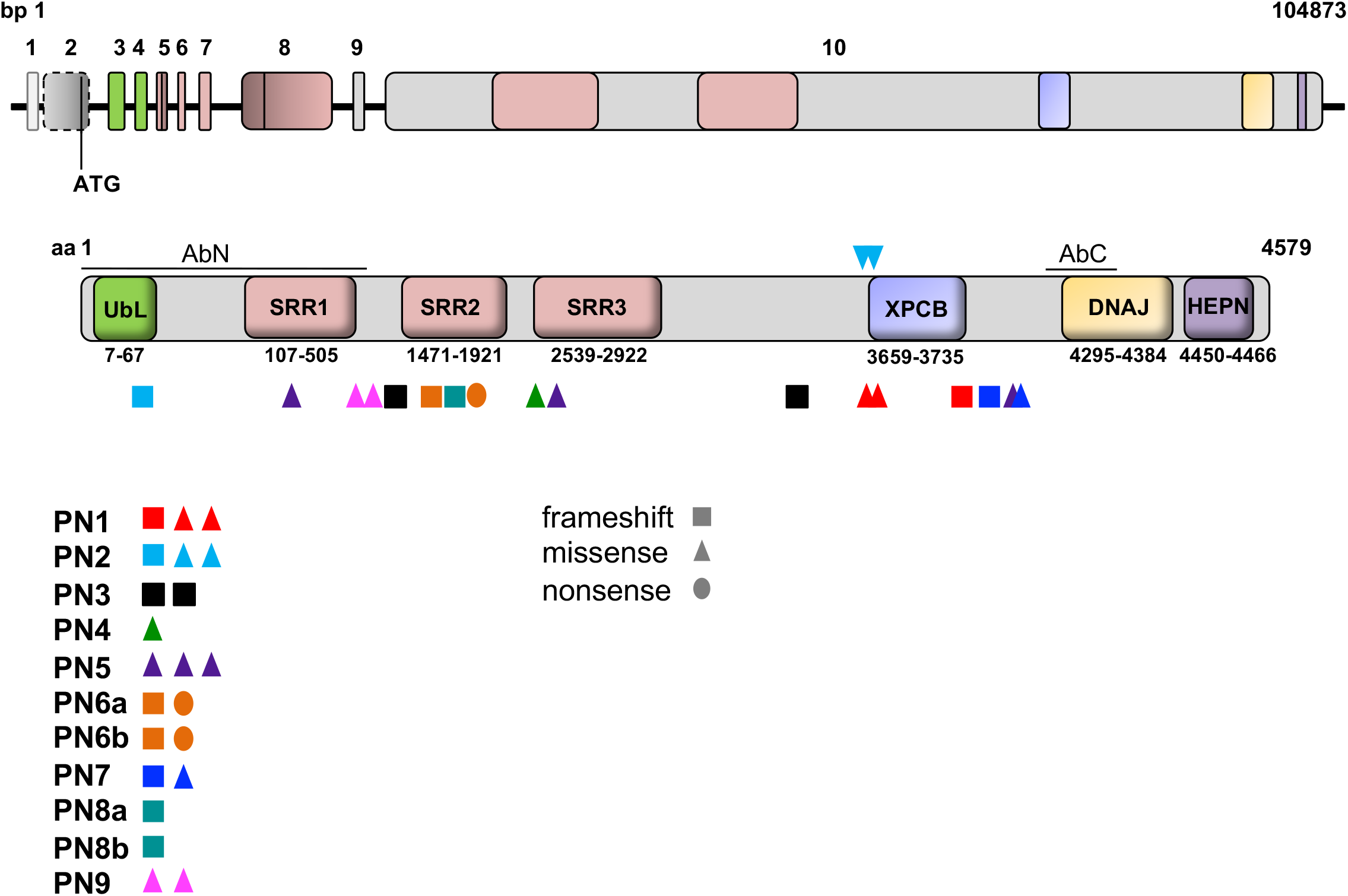
ARSACS patient fibroblasts analyzed in this study. Scheme of *SACS* gene and sacsin protein illustrating the identified functional domains and corresponding exons. For each patient (see **Table 1**), the type and position of the mutations on sacsin protein are indicated with a symbol and a color code. AbN and AbC antibodies used in this study are shown (bottom). Abbreviations: PN: patient; AbN/C: N/C-terminal anti-sacsin antibody; UbL: Ubiquitin-Like domain; SRR: Sacsin Repeating Region; XPCB: Xeroderma Pigmentosus group C protein Binding domain; HEPN: Higher Eukaryotes and Prokaryotes Nucleotide-binding domain; bp: base pair; aa: aminoacid.

Among the above-mentioned mutations, those carried by PN3 (p.[E723fs*15];[F3209fs*46]), PN6a and PN6b (p.[S1531fs*9];[R1645*]), PN8a and PN8b (p.[N1586fs*3];[deleted_allele]) and PN9 (p.[P536L];[R632W]) are reported in this study for the first time, at least in compound heterozygosity in the same patient. The other mutations are already published (PN1 and PN2 (Baets et al., 2010); PN4 (Terracciano et al., 2009); PN5 (Ricca et al., 2019); PN7 (Masciullo et al., 2012)) (**Fig. 1 and Table 1**).

To examine sacsin protein levels in the different ARSACS patients, we performed Western Blot (WB) using a commercially available C-terminal anti-sacsin antibody (AbC) (**Fig. 2A**) and a newly developed N-terminal anti-sacsin antibody (AbN) (**Fig. 2B**) (epitopes shown on the sacsin protein scheme in **Fig. 1**). AbN was designed and produced by our group, as no sacsin N-terminal antibodies efficiently recognizing sacsin in WB are commercially available.

**Figure 2.**
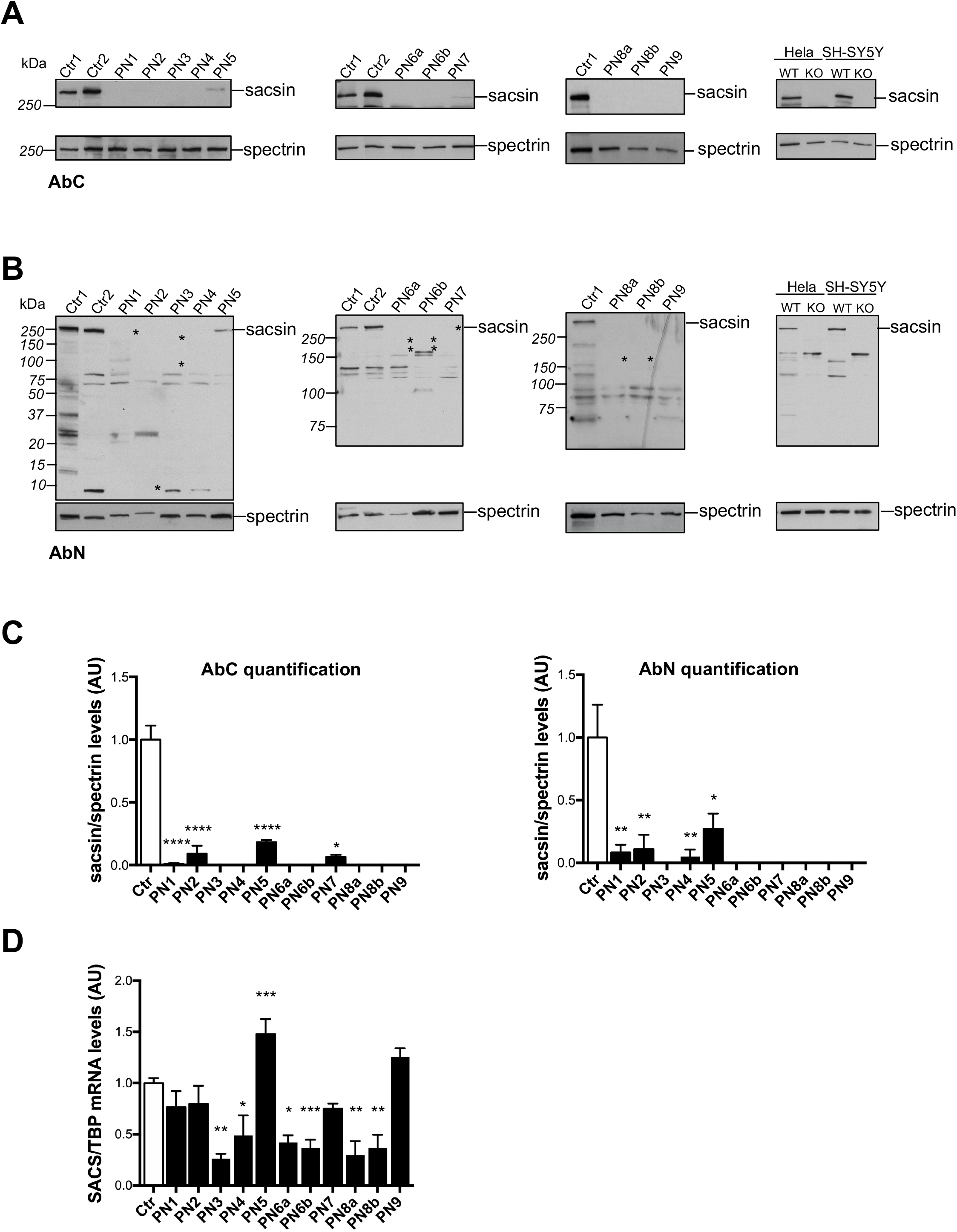
Sacsin protein is drastically reduced in ARSACS fibroblasts carrying different *SACS* mutations. (**A**) Representative WB showing residual sacsin protein in ARSACS patient-derived primary fibroblasts versus controls by using a C-terminal anti-sacsin antibody (AbC). Spectrin is used as loading normalizer. (**B**) Representative WB showing residual sacsin protein in ARSACS patient-derived primary fibroblasts versus controls by using AbN. Asterisks indicate the expected truncated sacsin protein: PN1=412 kDa; PN2=8 KDa; PN3=81kDa,358 kDa; PN6a and PN6b=170 kDa,180 kDa; PN7=410 kDa; PN8a and PN8b=175 kDa. (**C**) Quantification of sacsin levels relative to spectrin in WB experiments by using AbC and AbN. (**D**) *SACS* mRNA quantification by qRT-PCR in ARSACS patient-derived primary fibroblasts versus controls, normalized on *TBP* mRNA levels. In (**C-D**), data are presented as mean ± SEM. *p≤0.05; **p<0.01; ***p<0.001, ****p<0.0001 (unpaired-two tailed Student’s t-test). Abbreviation: Ctr = control.

To validate the specificity of AbN, we used sacsin knockout (KO) HeLa and SH-SY5Y cells we engineered by CRISPR-Cas9 technology (right panel, **Fig. 2B**).

Strikingly, with both antibodies we found that full-length sacsin protein is dramatically reduced or totally absent in all ARSACS patients, regardless of the nature of *SACS* mutations (**Fig. 2A and 2C**). This finding was unexpected for patients carrying monoallelic missense mutations, and even more for those carrying biallelic missense mutations.

We then employed the AbN to identify putative truncated products in patients carrying frameshift mutations on *SACS* gene. AbN recognizes a large epitope spanning the first sacsin methionine residue, till the end of exon 9 (residue 1-728). This epitope contains the sacsin UbL domain, a fairly common domain among proteins predicted to interact with the proteasome (Parfitt et al., 2009). AbN indeed recognizes other proteins carrying UbL domains (bands shared by controls and PNs in the WB in **Fig. 2B**). Considering the position of the premature stop codon for each mutation, the expected molecular weights of the sacsin fragments are reported in the legend and highlighted as asterisks in AbN WBs in **Fig. 2B**. Despite the AbN efficiently recognizes full-length sacsin protein, we were not able to appreciate any truncated sacsin fragments in WB by using AbN in any of the patients carrying frameshift mutations (**Fig. 2B**).

### *SACS* mRNA is reduced in ARSACS fibroblasts carrying frameshift mutations, while it is stable or even increased in those carrying missense mutations

To understand if the absence of full-length sacsin (and of truncated sacsin products) in ARSACS fibroblasts was due to mRNA instability, we first analyzed *SACS* mRNA levels by real-time PCR (qRT-PCR).

In ARSACS patients carrying only truncating mutations, *SACS* mRNA was evidently reduced compared to controls (PN3, PN8a, PN8b, PN6a and PN6b), suggesting a Non-sense Mediated Decay (NMD) of the mRNA. On the other hand, mRNA resulted stable (PN4, PN9) or even increased (PN5) in patients carrying missense mutations compared to controls. In ARSACS patients who are compound heterozygous for a missense and a frameshift mutation (PN1, PN2, PN7) we found no significant difference in the amount of *SACS* mRNA compared to controls, suggesting that the contribution of allele carrying the missense change determines the conserved amount of *SACS* mRNA (**Fig. 2D**).

### Inhibition of degradative systems does not rescue sacsin in ARSACS patients carrying missense mutations

Since *SACS* mRNA is stable and sacsin protein is dramatically reduced in ARSACS patients carrying missense mutations, we checked if mutant sacsin could undergo a faster post-translational degradation. Unfolded or aberrant proteins are usually targeted by the proteasome system or the autophagic system, the main molecular pathways involved in protein quality control and maintenance of cellular proteostasis (Morimoto and Cuervo, 2014). To test this hypothesis, we blocked cellular degradative systems by treating patient cells either with a proteasome inhibitor (MG-132, 1 μM, 24 or 3 hours) or an autophagy inhibitor (Chloroquine (CQ), 20 μM, 24 hours) (**Fig. 3A and 3C**). Both treatments did not rescue mutant sacsin in ARSACS patient fibroblasts carrying missense mutations. By blocking the proteasomal pathway for 24 hours, a statistically significant reduction of wild-type sacsin levels was observed in controls (**Fig. 3A,** graph). This suggested us a possible inhibition of protein synthesis due to a prolonged MG-132 treatment and consequent cellular stress induction (Gandin et al., 2010). We thus repeated the blockade of the proteasome reducing the time of treatment to 3 hours. In this condition, sacsin was stable in controls, but again not rescued in patients (**Fig. 3B**). We also blocked proteasome and autophagy together, but also in this case we did not see any increase in the amount of sacsin protein in patients (**Fig. 3D**).

**Figure 3.**
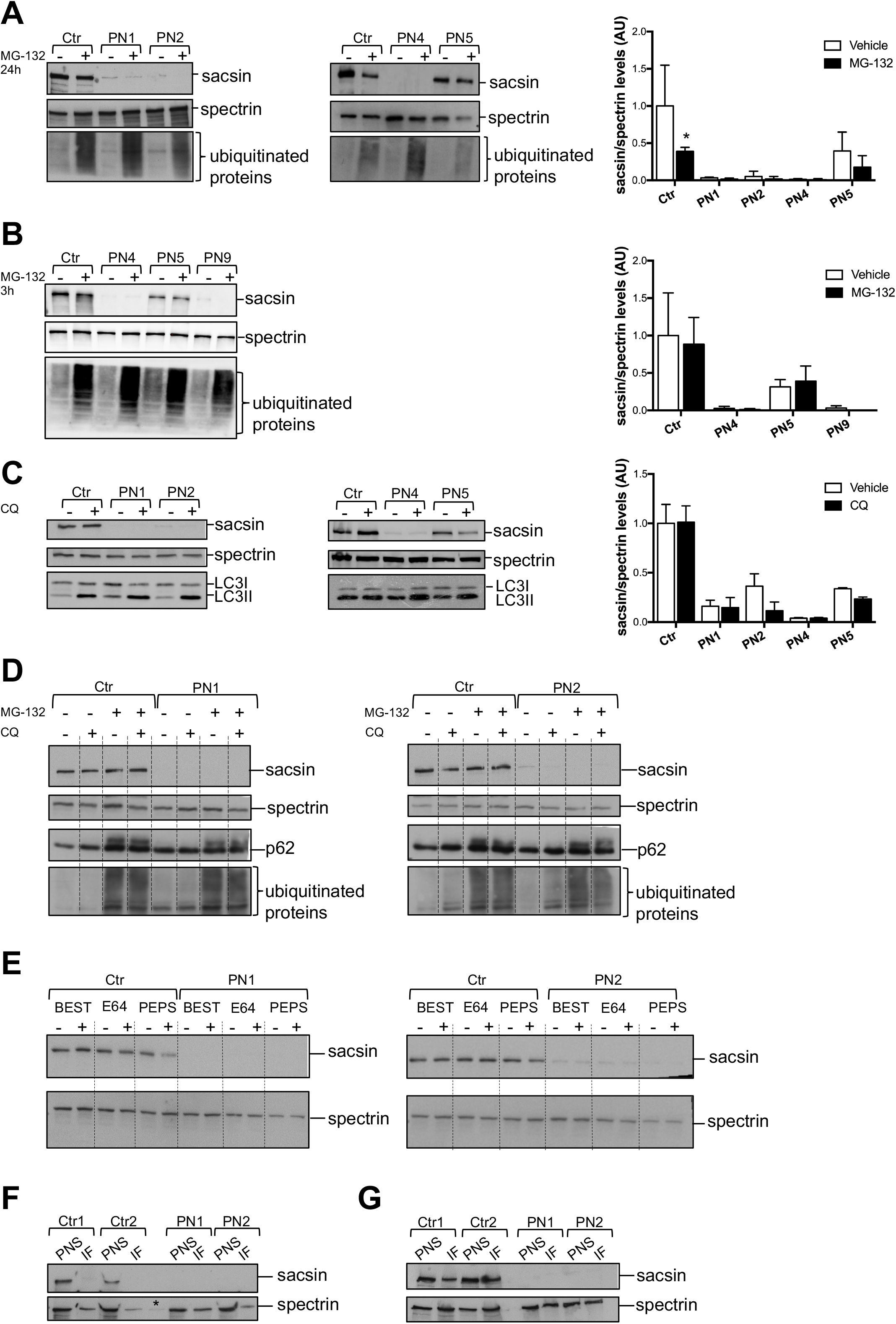
Mutant sacsin is not post-translationally degraded or aggregated. (**A**) Representative WB and quantification of sacsin levels in patient and control fibroblasts after proteasome blockade experiments, by using 1 μM MG-132 for 24 hours. Ubiquitinated proteins are used as readout of the treatment and spectrin as loading normalizer. (**B**) Representative WB and quantification of sacsin levels in patient and control fibroblasts after proteasome blockade experiments, by using 1 μM MG-132 for 3 hours. Ubiquitinated proteins are used as readout of the treatment. (**C**) Representative WB and quantification of sacsin levels in patient and control fibroblasts after autophagy blockade experiments, by using 20 μM Chloroquine (CQ) for 24 hours. LC3I conversion in LC3II is used as readout of the treatment. (**D**) Representative WB of sacsin levels in patient and control fibroblasts upon proteasome plus autophagy blockade, by using 0.5 μM MG-132 and 10 μM CQ for 18 hours. P62 is used as readout of the CQ treatment and ubiquitinated proteins of the MG-132 treatment. (**E**) Representative WB of sacsin levels in patient and control fibroblasts upon protease inhibition by using different protease inhibitors for 48 hours (repeating the treatment at 24 hours). (**F**) Representative WB of sacsin levels in patient and control soluble and insoluble fractions, obtained by pellet sonication. Asterisk represents a leak of material from the adjacent well. (**G**) Representative WB of sacsin levels in patient and control soluble and insoluble fractions resuspended in Laemmli sample buffer. In (**A–C**), data are presented as mean ± SEM. n=at least 3, *p≤0.05 (unpaired-two tailed Student’s t-test). Abbreviations: + = treated; − = vehicle; BEST= bestatin; PEPS= pepstatin; PNS= post-nuclear surnatant; IF= insoluble fraction.

We then investigated if sacsin could be degraded by specific proteases. We inhibited cysteine proteases with E64, amino peptidases with bestatin and aspartyl proteases with pepstatin. The low amount of mutant sacsin in ARSACS patients was not modified by any of these treatments (**Fig. 3E**). We took in consideration that sacsin carrying missense mutations could be completely translated, but undetectable by standard WB procedures due to its misfolding and/or aggregation. To address this hypothesis, we performed different experiments to solve aggregates in biochemical assays. We checked for putative mutant sacsin aggregates by loading in SDS-PAGE both the soluble and the insoluble fractions of control and patient fibroblasts (PN1, PN2) obtained by pellet sonication (**Fig. 3F**) or by direct resuspension in Laemmli sample buffer (**Fig. 3G**). We were unable to detect sacsin aggregates in the insoluble fractions in patient cells. We also performed WB using mixed acrylamide-agarose gel to solve aggregates without rescuing mutant sacsin levels (not shown).

Altogether, these results indicate that the drastic reduction of mutant sacsin in ARSACS patients is not caused either by a faster post-translational degradation of the protein or aggregation.

### Translation of mutant sacsin carrying missense mutations is not blocked in ARSACS patients

At this point we considered that the absence of sacsin carrying missense mutations could be due to its inefficient translation. We first excluded a general problem in translation, as polysomal profiles showed a similar pattern in ARSACS patients and controls (**Fig. 4A**). In order to know if wild-type *SACS* mRNA could be subjected to a translational regulation *per se*, we performed meta-analysis of ribosome profiling data downloaded and analyzed exploiting the GWIPS-viz platform (Michel et al., 2014). This analysis showed little accumulation of ribosomes in the 5’UTR of *SACS* mRNA, and a homogenous density of ribosomes across the 13737 nucleotide long mRNA sequence, suggesting the absence of strong regulatory sequences in the 5’UTR (**Fig. 4B**). To specifically assess the translation of sacsin, we investigated if *SACS* mRNA carrying missense mutations was associated to intact ribosomes (polysomes) or split ribosomes (monosomes, due to a blockade of translation). We thus performed qRT-PCR for sacsin on RNA extracted from the different ribosomal fractions (monosomes, light polysomes, heavy polysomes) for ARSACS patients and controls. We found that mutant *SACS* mRNA was associated to the polysomal fractions (the actively translating fractions) in ARSACS patients (PN4, PN5, PN9) as well as in controls (**Fig. 4C**, left), demonstrating that there is no blockade of its translation. The same mRNA distribution on the different ribosomal fractions was observed for a housekeeping gene (*TBP*) tested as control (**Fig. 4C**, right).

**Figure 4.**
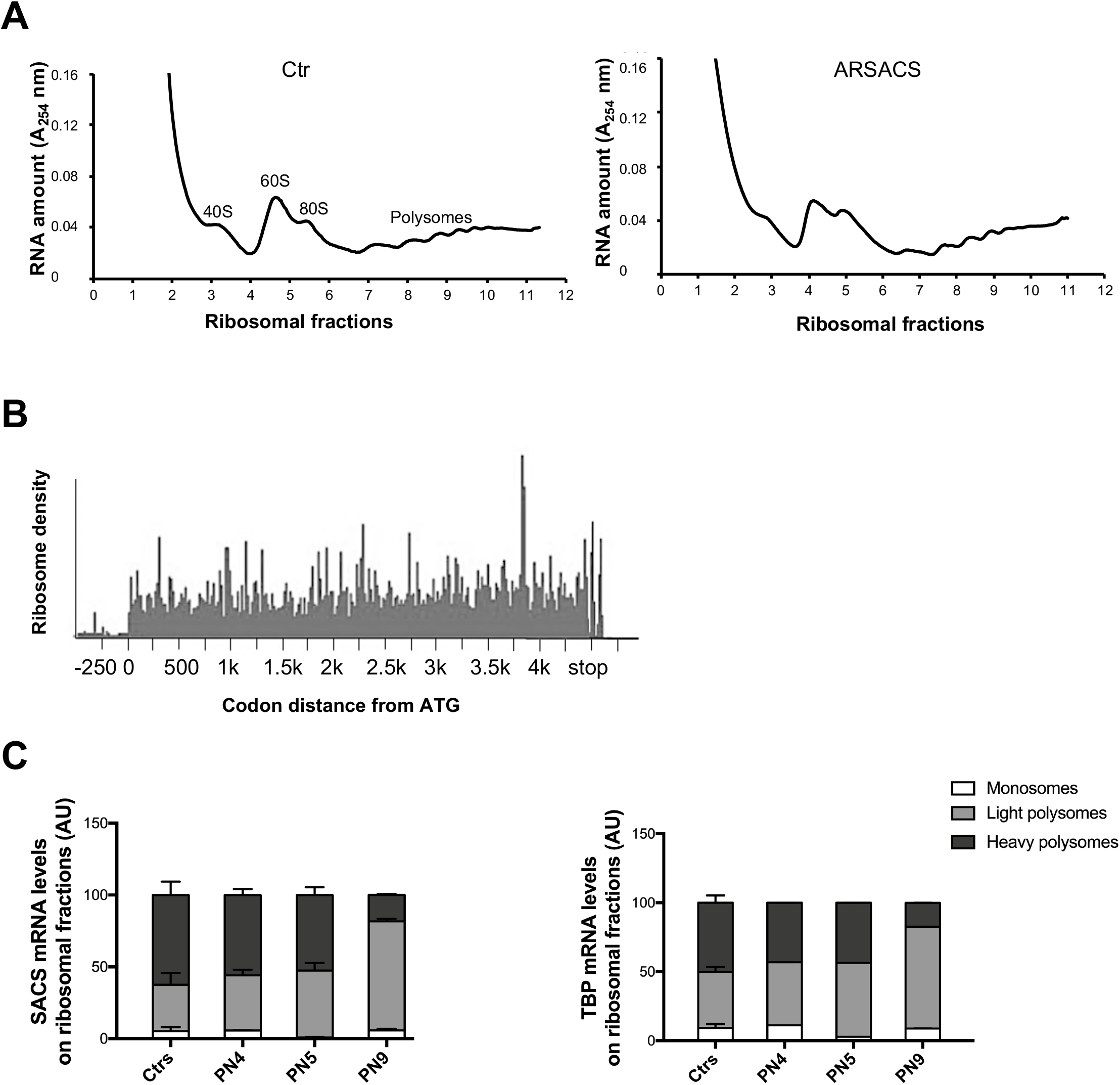
SACS mRNA carrying missense mutations is associated to the polysomal fractions. (**A**) Representative ribosomal profile graphs in control (left) and ARSACS patient fibroblasts (right). (**B**) Ribosomal density on the human sacsin mRNA, retrieved and analyzed from available riboseq datasets. (**C**) qRT-PCR data of the *SACS* mRNA distribution in different ribosomal fractions: monosomes, light and heavy polysomes in ARSACS patient fibroblasts (PN4, PN5, PN9) compared to 3 different controls (left). *TBP* mRNA distribution is shown as housekeeping gene analyzed in the same cells (right).

### Nascent mutant sacsin products are cotranslationally ubiquitinated and degraded

At this point, having experimentally excluded all the possible molecular mechanisms accounting for sacsin reduction in ARSACS patients, we considered that mutant sacsin could be degraded during translation. Two different quality control (QC) mechanisms exist during translation, both leading to the ubiquitination and degradation of nascent proteins. The first one, the ribosomal QC, is associated to the mRNA degradation and ribosome stalling/splitting (i.e the NMD, the No-Go Decay etc.) (Inada, 2017). The other one, the cotranslational QC, senses the correct folding of the nascent chain, promoting the degradation of the protein while it is being translated, and occurs in the presence of a stable mRNA associated to intact ribosomes (Wang et al., 2015). This second scenario is the most plausible in the case of mutant sacsin carrying missense mutations, since we did not detect any reduction of the mRNA. Although poorly characterized, cotranslational QC is predicted to occur for big and multimodular proteins, whose folding takes place cotranslationally and proceeds in a domain-wise manner (Liutkute et al., 2020). The players involved in nascent protein degradation associated to cotranslational QC are completely unknown so far.

As formal proof of the cotranslational degradation of sacsin carrying missense mutations, we looked for the presence of ubiquitinated degradation products of nascent mutant sacsin in ARSACS patient cells carrying missense mutations only. Thus, we immunoprecipitated sacsin with AbN (in the view of recognizing N-terminal sacsin fragments of unpredictable molecular weights) in ARSACS patients (PN5, PN9) versus controls. Since putative mutant sacsin intermediates should be ubiquitinated and degraded by the proteasome, before the immunoprecipitation (IP) we treated cells with the proteasome inhibitor MG-132 (1uM, 3 hours) to improve their detection. We efficiently immunoprecipitated full-length sacsin with AbN in control and (to a lower extent as expected) in patient cells (**Fig. 5A**). In ARSACS fibroblasts, sacsin degradation products are detectable especially in the higher part of the gel, in particular with the anti-ubiquitin immunodecoration (**Fig. 5A,** right). To better resolve the higher part of the gel we decided to reload an independent sacsin IP in a 6% gel (**Fig. 5B**). Sacsin degradation products were clearly visible in the IP lanes of ARSACS patients, especially when immunodecorated with the AbN (**Fig. 5B**, left). To enhance sacsin ubiquitination, we transfected control and patient cells with Ubiquitin-HA construct and immunoprecipitated sacsin in the same conditions as before. We reconfirmed the detection of sacsin degradation products in ARSACS patients (**Fig. 5C**) and, interestingly, we noticed that the differential bands present in the IP samples of patients were more evident by immunodecoration with anti-HA (revealing the ubiquitinated sacsin products) (**Fig. 5C,** right).

**Figure 5.**
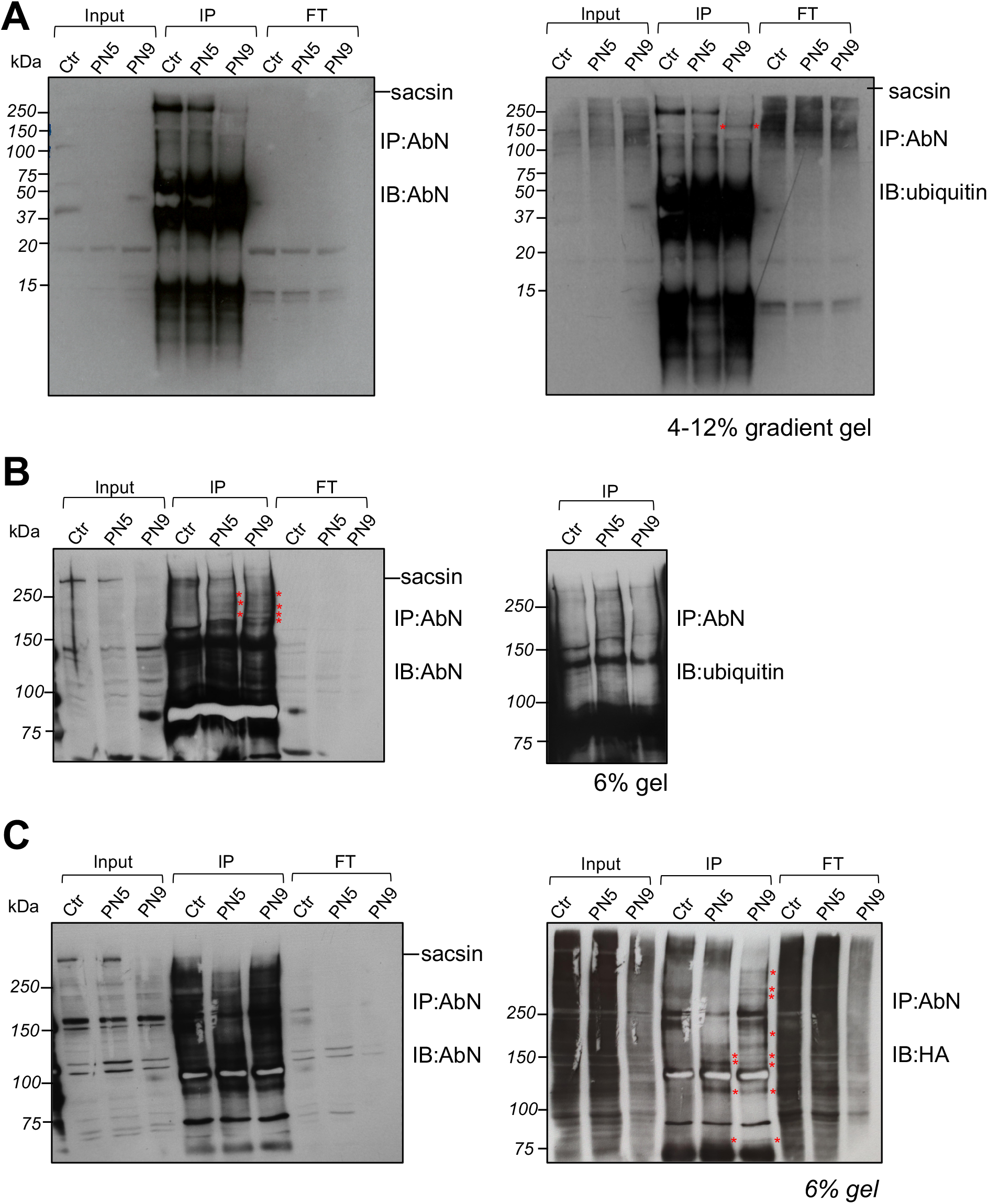
Sacsin protein carrying missense mutations is co-translationally ubiquitinated and degraded. (**A**) Sacsin IP by using AbN in ARSACS patient fibroblasts carrying missense mutations (PN5, PN9) and a control treated with 1 μM MG-132 for 3 hours. 4-12 % gradient gel showing proteins from high molecular weights to 15 kDa, immunodecorated with AbN (left) or ubiquitin (right). (**B**) Independent experiment conducted in the same conditions loaded onto a 6% gel to resolve putative differential N-terminal sacsin bands in patients compared to control. Immunodecoration with AbN (left WB) or ubiquitin (right WB, red asterisks indicate N-terminal sacsin products visible only in the patient IP samples). (**C**) Sacsin IP by using AbN in ARSACS patient fibroblasts carrying missense mutations (PN5, PN9) and a control upon Ubiquitin-HA overexpression, treated with 1 μM MG-132 for 3 hours. 6 % gel showing immunodecoration with AbN (left) and with anti-HA revealing ubiquitinated proteins (right). Red asterisks indicate differential N-terminal sacsin products visible only in the patient IP samples. Abbreviations: IP: immunoprecipitation; FT: Flow through fractions; IB: immunoblotting.

According to our results, we can arrange a final model of the first event of ARSACS pathogenesis: in the presence of frameshift or nonsense mutations, *SACS* mRNA is degraded; in the presence of missense mutations, sacsin fails to fold and undergoes cotranslational degradation (**Fig. 6**). In both cases the result is the absence/striking reduction of sacsin.

**Figure 6.**
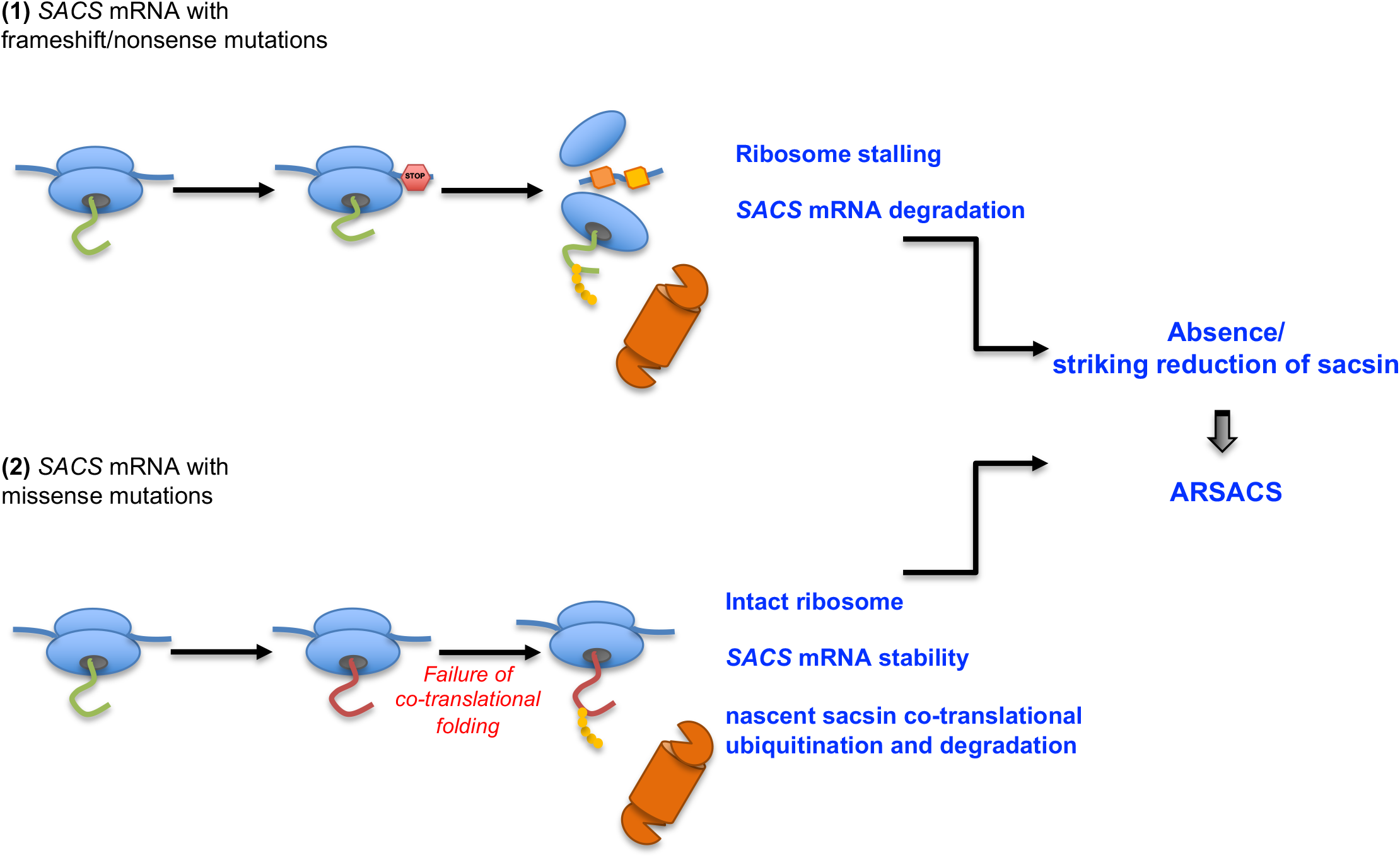
Full length sacsin protein is almost absent in ARSACS patients, independently of the nature of the mutations. Final model explaining why mutant sacsin is almost absent in patient cells. In (1) sacsin is prevented to be translated because the *SACS* mRNA carrying frameshift mutations is degraded by nucleases, while in (2) the *SACS* mRNA carrying missense mutations is stable and sacsin is cotranslationally degraded. The proteasome is the degradative system implied in these pathways (drawn in orange). Green nascent chain: before sensing the mutation; red nascent chain: after sensing domain misfolding.

## Discussion

In this work, we studied in detail for the first time the effects of different ARSACS-causing mutations on sacsin stability, at both mRNA and protein level. The cohort of ARSACS patients that we analyzed encompasses a wide range of diverse types of *SACS* mutations localized in different regions of the gene. Our results demonstrate that sacsin is barely detectable in ARSACS patients regardless of the nature or position of the mutation along the gene. While this result was expected for patients carrying biallelic truncating mutations (whose mRNA is unstable and degraded), it was not anticipated for patients carrying missense mutations, especially on both alleles. We indeed discovered that sacsin carrying missense mutations in any site of its sequence is cotranslationally ubiquitinated and degraded, rarely reaching its mature size. We surmise that this mechanism prevents the even more dangerous production of a huge misfolded protein, potentially highly prone to aggregation in the crowded cytosolic environment.

Since ARSACS presents variability in its clinical presentation, efforts have been dedicated to find a genotype-phenotype correlation, to better tailor a precision medicine approach, in terms of disease prognosis, family counselling and for future clinical trials. Our data however do not confirm previous reports only based on *in silico* analyses correlating the nature and/or position of the *SACS* mutations with the severity of the disease (Romano et al., 2013; Xiromerisiou et al., 2020). Here, we demonstrated that loss of function in ARSACS is caused by loss/striking reduction of sacsin, independently of the mutation.

Excluding the Quebec cohort, where the original c8844delT *SACS* mutation is highly prevalent, the 2020 worldwide ARSACS scenario shows that most mutations are missense, followed by small deletions, frameshift, nonsense and small insertions (Xiromerisiou et al., 2020). In our cohort of compound heterozygous patients, two of them carry biallelic missense mutations and five of them monoallelic missense, for a total of nine different missense changes. In all cases, the full-length protein is almost absent in patients with missense mutations (hitting different regions of the protein, from the N-terminus to the C-terminus) as well as in patients with truncating mutations. Only in PN5 a residue of sacsin was detected with both AbC and AbN antibodies, which is anyway about 20% of controls, whereas in PN9 (who carries biallelic missense as well) sacsin is not detectable.

Our findings are supported by the recently published of *Sacs^R272C/R272C^* knockin mouse model, carrying a human pathogenetic mutation in the UbL domain (Lariviere et al., 2019). This model presents a barely detectable sacsin protein despite a stable mRNA and, accordingly, a phenotype overlapping with the one of the *Sacs* knockout mouse (Lariviere et al., 2015; Lariviere et al., 2019), confirming the conservation of the cotranslational degradation of mutant sacsin also in cerebellum. In addition, a striking reduction of sacsin was shown in other two reports of ARSACS mutations in patient fibroblasts (Duncan et al., 2017; Thiffault et al., 2013), with no further mechanistic insights.

We found that *SACS* mRNA was stable or even increased also in the ARSACS compound heterozygous patients carrying biallelic or monoallelic missense mutations, excluding mRNA degradation as the possible explanation for the absence of sacsin. The failure in rescuing full-length sacsin upon the inhibition of all cellular degradative systems and several proteases excluded faster post-translational degradation. Also, mRNA of sacsin turned out to be associated with polysomes in ARSACS patients as well as in controls, rejecting the hypothesis of defective translation. We instead discovered that sacsin protein carrying missense mutations is cotranslationally ubiquitinated and degraded.

To preserve proteostasis, eukaryotic cells must not only promote accurate folding, but also prevent the accumulation of misfolded species that may arise from inefficient folding, errors in translation, and aberrant mRNAs. A growing body of evidence indicates that large and multimodular proteins start folding cotranslationally. For such proteins, domain-wise cotranslational folding helps reaching their native states (which would be otherwise highly challenging in the overcrowded cytosol) and may reduce the probability for off-pathways and aggregation-prone conformations (Liutkute et al., 2020; Pechmann et al., 2013; Waudby et al., 2019). Also, many studies indicate that a sizable portion of nascent chains are cotranslationally ubiquitinated and degraded by the proteasome in physiological conditions, both in yeast (Duttler et al., 2013; Willmund et al., 2013) and in mammalian cells (Wang et al., 2013). This likely applies to wild-type sacsin, a 520 kDa protein with a complex multimodular architecture containing repeated motifs, whose folding may take minutes to hours to complete. We hypothesize that cotranslational folding and degradation occur physiologically for sacsin as mechanisms of QC. In the case of sacsin carrying a missense mutation, after the unsuccessful attempts of chaperones to fold the nascent chain, the latter is constitutively ubiquitinated and degraded, preventing the synthesis of a misfolded full-length protein. This compartmentalized surveillance mechanism results in a loss-of-function, avoiding a potentially more dangerous toxic gain-of-function of mutant sacsin in the cytosol, which may promiscuously interact with non-specific proteins forming harmful aggregates.

We provided the formal proof of cotranslational degradation of mutant sacsin with the IP with AbN in condition of proteasome inhibition, which revealed the presence of several specific ubiquitinated sacsin products at lower molecular weight compared to the full-length in ARSACS patients carrying missense mutations.

The low residual amount of mutant sacsin that is present in some patients (PN5 and PN7), even if barely detectable, may be due to a quote of mutant protein escaping from cotranslational degradation, because C-terminally localized mutations are presumably more permissive. Indeed, the stability and susceptibility of different folding domains to degradation may vary depending on the polypeptide sequence (Wolff et al., 2014).

Other studies observed cotranslational ubiquitination of cystic fibrosis transmembrane conductance regulator (CFTR) carrying the ΔF508 mutation (Sato et al., 1998), wild-type apolipoprotein B100 (Zhou et al., 1998), and homeodomain interacting protein kinase 2 (HIPK2) carrying artificial mutations (Muller et al., 2021) by either *in vitro* translation or in overexpression conditions. To our knowledge this is the first report showing *in vivo* in endogenous conditions the cotranslational degradation of a cytosolic protein (carrying any type of missense mutation) as the cause of a human disease.

Our data identifies lack of sacsin protein as unifying mechanism shared by different *SACS* mutations, with multiple important implications for ARSACS disease managing. For the diagnosis, the evaluation of sacsin levels could be included in the clinical genetics practice to establish a definite ARSACS diagnosis, or even as pre-screening in high probable cases to avoid expensive next-generation sequencing panel analysis, and certainly to validate variants of uncertain clinical significance (VUS). For designing therapeutic strategies, cotranslational degradation of mutant sacsin makes unproductive any post translational approach. Further studies are of course needed to explain the pronounced intra- and interfamily variability of ARSACS patients, such as the impact of gene modifiers, epigenetic factors and environmental differences on disease phenotype.

## Materials and Methods

### Patient consent

Subjects’ consent was obtained according to the Declaration of Helsinki and was approved by the local ethical committee of Ospedale San Raffaele, IRCCS Stella Maris Pisa and the Antwerp University Hospital.

### Human primary fibroblast derivation from skin biopsies and cell culture

Patient-derived skin fibroblasts were obtained following standard protocols during diagnostic procedures. Fibroblasts were cultured in D-MEM medium supplemented with 20% FBS, 1mM sodium pyruvate, 2mM L-glutamine and 100 U/ml penicillin-streptomycin. All cell culture reagents were from Thermo Fisher Scientific (Waltham, MA, USA).

### Human primary skin fibroblasts immortalization

For polysomal profiling and IP experiments we used immortalized fibroblasts. Cells were transduced with a lentiviral vector, kindly gifted by Mario Squadrito’s lab (1:2500; initial concentration 6*10^9^ TU/mL) carrying SV-40 gene under the spleen focus forming virus (SFFV) promoter. Growth of cells was observed and compared to non-transduced human primary skin fibroblasts.

### Antibodies, drugs and reagents

Commercially available antibodies were used in Western blots (WB), for the detection of sacsin (Anti-sacsin AbC ab181190. Abcam, Cambridge, UK), spectrin (Anti-spectrin MAB1622 Merck Millipore, Burlington, MA, USA), ubiquitinated proteins (Anti-ubiquitin ab134953, Abcam), p62 (Anti-p62/SQSTM1 P0067 Merck Millipore), LC3I-II (Anti-LC3A/B ab58610. Abcam), HA (anti-HA Epitope Tag 16b12, Biolegend, San Diego, CA, USA). Secondary antibodies included HRP-conjugated anti-mouse and anti-rabbit IgG (Amersham Bioscience, Buckinghamshire, UK).

*In vitro* treatments were carried out with different compounds: 1 μM MG-132 24 hours or 3 hours (Merck Millipore), 20 μM Chloroquine (CQ) 24 hours (Merck Millipore) or together (0.25 μM MG-132 + 10 μM CQ, 18 hours); protease inhibitors (5 μM E64, 10 μM Bestatin, 5 μM Pepstatin, 48 hours (Merck Millipore)). After the treatments, cells were harvested and lysed for biochemical assays.

### Cell lysis, SDS-PAGE and WB analyses

Cells were lysed as previously described (Longo et al., 2020). Protein quantification was performed with Bradford assay accordingly to the manufacturer’s instructions. Samples were resuspended in SDS sample buffer (5.8 mM TrisHCl pH 6.8, 5% glycerol, 1. 6% SDS, 0.1 M DTT, 0.002% bromophenol blue), boiled and loaded onto SDS-PAGE followed by immunoblot.

### Anti N-terminal sacsin antibody (AbN) generation

We generated a new rabbit polyclonal anti-sacsin antibody recognizing the N-terminal portion of sacsin (ID Q9NZJ4 (SACS_HUMAN)). We decided to develop a new anti-sacsin antibody (AbN) raised against a polypeptidic region of sacsin including aminoacids from 1 to 728 (all the exons till the giant exon 10). The antibody was produced by the antibody production service of Biomatik (Cambridge, Ontario, Canada).

### Protein aggregate detection

Cells were lysed by using Dounce homogenizer in 100 mM Tris-HCl pH 7.4, 150 mM NaCl, 1 mM EDTA pH 8 + protease inhibitor cocktail (Merck Millipore), centrifugated 1000g x 10 minutes. Post-nuclear supernatant (PNS) was collected, and pellets were resuspended in 500 μL lysis buffer and sonicated (20 seconds pulse-10 seconds stop; repeated twice) or directly resuspended in SDS sample buffer. PNS and pellet were quantified and equal amount were loaded on SDS-PAGE.

### *SACS* deletion by CRISPR/Cas9

Traditional 20 bp-NGG spCas9 gRNAs targeting *SACS* coding region were designed with CHOPCHOP web tool. The gRNA sequences were: gRNA-1 5’ TGCTCCTGCGGTTATCAGTA 3’; gRNA-2 5’ GTAGGCCATGCAATTCTCAT 3’. Oligos encoding the gRNAs were annealed and cloned into pCas-Guide-EF1a-GFP CRISPR/Cas9 plasmid (OriGene), according to the manufacturer instructions. To delete *SACS* gene, HeLa cells and SH-SY5Y cells were transfected with CRISPR/Cas9 plasmid carrying gRNA-1 or gRNA-2, with Metafectene PRO (Biontex) and Lipofectamine 3000 (ThermoFisher Scientific) respectively. Two days after transfection, GFP positive cells were sorted by FACS Aria Fusion (Becton Dickinson), and plated at single cell density in 96-multiwell plates. Monoclonal cell lines were expanded from wells with single colonies and sacsin KO was validated by immunoblot with AbC antibody and by Sanger sequencing of target genomic region. Primers for gRNA-1 target region were: forward primer 5’ AGCAAAAGGAGCAACGTCTG 3’; reverse primer 5’ GCTCTTTTCCATCTCCAGACG 3’. Primers for gRNA-2 target region were: forward primer 5’ AGCCAAAACCCTCTTACTGG 3’; reverse primer 5’ AGTGGCTCTCTTTGTCCTGA 3’.

### RNA extraction and Real Time-PCR (qRT- PCR)

Total mRNA from primary fibroblasts was extracted with Trizol Reagent following manufacturer’s instructions. RNA was reverse transcribed in cDNA using SuperScript IV Reverse Transcriptase (ThermoFisher Scientific) with random hexamers. For each reverse transcription experiment, *GAPDH* PCR was performed to check the cDNA and its absence in the non-reverse transcribed samples to avoid genomic DNA contamination (not shown). *GAPDH* primers: forward primer 5’ CCACCCAGAAGACTGTGGAT 3’; reverse primer 5’ GTTGAAGTCAGAGGAGACCACC 3’. qRT- PCR was performed based on SYBR Green chemistry (Light Cycler 480. SYBR Green I master, Roche). *SACS* primers for the analysis of total *SACS* mRNA levels in patients and controls were: forward primer 5’ TTTTCAGTTGCGAGGGGTTG 3’; reverse primer 5’ TCCTGGCTTGGGAGGTAAAG 3’. To normalize *SACS* mRNA levels, we used TATA Binding Protein (*TBP*) mRNA levels. *TBP* forward primer 5’ ACGCCGAATATAATCCCAAG 3’; reverse primer 5’ GCACACCATTTTCCCAGAAC 3’.

### Polysomal profile analysis

Immortalized growing fibroblasts were treated with 100 μg/mL cycloheximide for 15 minutes and lysed on ice in 50 mM TrisHCl pH 7.5, 100mM NaCl, 30 mM MgCl_2_, 0.1% NP-40, 100 μg/mL cycloheximide, 40 U/mL RNAsin (Promega) and protease inhibitor cocktail (Merck Millipore). After centrifugation at 14000 g for 10 minutes at 4°C, the supernatants with equal amounts of RNA were loaded on a 15–50% sucrose gradient and centrifuged at 4°C in a SW41Ti Beckman rotor for 3 hours 30 minutes at 39000 rpm. The gradients were analyzed by continuous flow absorbance at 254 nm, recorded by BioLogic LP software (Bio-Rad), and all fractions were collected for the subsequent examination of translated mRNAs. Briefly, these fractions were divided into monosomes, (from the top of the gradient to the 80S peak), light polysomes and heavy polysomes. Samples were incubated with proteinase K and 1% SDS for 1 hour at 37°C. RNA was extracted by phenol/chloroform/isoamyl alcohol method. After treatment of RNA with RQ1 RNase-free DNase (Promega), reverse transcription was performed according to SuperScript III First-Strand Synthesis kit instructions (Thermo Fisher Scientific). Complementary cDNA (100 ng) was amplified with the appropriate SYBR green specific primers. To analyze *SACS* mRNA, different specific primers were used: forward primer 5’ GGCAATTTTGTCCCCTTCTCC 3’ reverse primer 5’ GGTCTTCCTCGGGTTTGGG 3’. qRT-PCR for *TBP* (as housekeeping gene) was used as control of translation. For the analysis of targets from monosome and polysome fractions, the data are quantified as the percentage of expression in each fraction.

### Immunoprecipitation of N-terminal sacsin fragments

Immortalized fibroblasts (transfected for 16 hours with Ubiquitin-HA plasmid or not) were plated in T75 flasks (4.5 x 10^6^ cells) and after 24 hours were treated for 3 hours with 1 μM MG-132 and harvested. Cells were lysed in RIPA buffer, then incubated for 30 minutes on ice and centrifuged at 8000 g for 10 minutes. Supernatant was collected and quantified by Bradford standard procedures. The lysate was incubated for 2 hours in a rotating wheel at 4°C with Dynabeads Protein A (#10002D Dynabeads Protein A, ThermoFisher Scientific) as a preclearing step. A different aliquot of Dynabeads Protein A was incubated with AbN (6 μg of antibody per 700 μg of protein) for 30 minutes at room temperature and then washed in RIPA buffer. Precleared lysates were then incubated overnight at 4°C on a wheel with Dynabeads Protein A-AbN. The day after, unbound proteins were collected as flow through (FT) while the immunoprecipitated proteins (IP) were washed 5 times in RIPA buffer and then eluted in strong elution buffer (8 M Urea, 100 mM Tris HCl pH 8), followed by separation onto SDS-PAGE. Ubiquitin-HA plasmid was kindly provided by S. Polo’s lab, IFOM, Milan.

### Statistical analyses

Continuous variables were summarized by their mean values and Standard Error of the Mean (SEM). Differences in protein levels between patients and controls were assessed by densitometric analysis of WB bands from at least three independent experiments using Image J, followed by student *t*-test analysis.

## Acknowledgements

We thank all patients for collaborating with this study. We thank Ignazio Lopez for deriving skin biopsies from patients. We are grateful to Roberto Sitia for critical discussion.

This project was supported by the Italian Ministry of Health # RF-2016-02361610 and Ataxia Charlévoix-Saguenay Foundation (FM). FL was recipient of a Fondazione Centro San Raffaele- Fronzaroli fellowship. This work was supported by the Association Belge contre les Maladies Neuromusculaire (ABMM) and the EU Horizon 2020 program (Solve-RD, No 779257). JB is supported by a Senior Clinical Researcher mandate of the Research Fund - Flanders (FWO) under grant agreement number 1805021N. JB is a member of the μNEURO Research Centre of Excellence of the University of Antwerp and of European Reference Network for Rare Neuromuscular Diseases (ERN EURO-NMD).

## Author contributions

FM, SB and FL conceived the study, interpreted the results and wrote the manuscript. FL, DDR, AM, DF performed the experiments, analyzed data and helped writing the manuscript. JB, MS and FS are patient neurologists and provided skin biopsies. All authors critically discussed the data.

## The paper explained

### PROBLEM

ARSACS is a neurodegenerative disorder characterized by cerebellar ataxia, spasticity and peripheral neuropathy. It is caused by mutations in the *SACS* gene encoding sacsin, a huge multimodular protein of unknown function. More than 200 *SACS* mutations have been described worldwide, making ARSACS the second most common form of recessive ataxia after Friedreich’s ataxia. ARSACS phenotypic spectrum is highly variable among patients, in terms of severity and clinical presentation.

### RESULTS

In this study, we demonstrated that the clinical variability in ARSACS does not depend on the type/position of the *SACS* mutations. We indeed we found that sacsin is almost absent in a large set of ARSACS patient skin fibroblasts, regardless of the nature of the mutation. This is caused by a premature degradation of sacsin, which emerges as a novel cause of a human disease.

### IMPACT

Our data explain the mechanism underlying the lack of genotype-phenotype correlation in ARSACS. We propose that sacsin level should be evaluated to define ARSACS diagnosis. Moreover, our results make unproductive any translational approach targeting mutant sacsin.

## For more information

OMIM database:
ARSACS: https://www.omim.org/entry/270550
*SACS*: https://www.omim.org/entry/604490
www.hgmd.cf.ac.uk
https://www.lovd.nl
https://chopchop.cbu.uib.no
imagej.nih.gov/ij/index.html
https://gwips.ucc.ie/

## References

Anderson, J.F., Siller, E., Barral, J.M., 2010. The sacsin repeating region (SRR): a novel Hsp90-related supra-domain associated with neurodegeneration. J Mol Biol 400, 665–674.

Baets, J., Deconinck, T., Smets, K., Goossens, D., Van den Bergh, P., Dahan, K., Schmedding, E., Santens, P., Rasic, V.M., Van Damme, P., Robberecht, W., De Meirleir, L., Michielsens, B., Del-Favero, J., Jordanova, A., De Jonghe, P., 2010. Mutations in SACS cause atypical and late-onset forms of ARSACS. Neurology 75, 1181–1188.

Bouchard, J.P., Barbeau, A., Bouchard, R., Bouchard, R.W., 1978. Autosomal recessive spastic ataxia of Charlevoix-Saguenay. Can J Neurol Sci 5, 61–69.

Bouhlal, Y., Amouri, R., El Euch-Fayeche, G., Hentati, F., 2011. Autosomal recessive spastic ataxia of Charlevoix-Saguenay: an overview. Parkinsonism Relat Disord 17, 418–422.

Duncan, E.J., Lariviere, R., Bradshaw, T.Y., Longo, F., Sgarioto, N., Hayes, M.J., Romano, L.E.L., Nethisinghe, S., Giunti, P., Bruntraeger, M.B., Durham, H.D., Brais, B., Maltecca, F., Gentil, B.J., Chapple, J.P., 2017. Altered organization of the intermediate filament cytoskeleton and relocalization of proteostasis modulators in cells lacking the ataxia protein sacsin. Hum Mol Genet 26, 3130–3143.

Duttler, S., Pechmann, S., Frydman, J., 2013. Principles of cotranslational ubiquitination and quality control at the ribosome. Mol Cell 50, 379–393.

Engert, J.C., Berube, P., Mercier, J., Dore, C., Lepage, P., Ge, B., Bouchard, J.P., Mathieu, J., Melancon, S.B., Schalling, M., Lander, E.S., Morgan, K., Hudson, T.J., Richter, A., 2000. ARSACS, a spastic ataxia common in northeastern Quebec, is caused by mutations in a new gene encoding an 11.5-kb ORF. Nat Genet 24, 120–125.

Gagnon, C., Brais, B., Lessard, I., Lavoie, C., Cote, I., Mathieu, J., 2018. From motor performance to participation: a quantitative descriptive study in adults with autosomal recessive spastic ataxia of Charlevoix-Saguenay. Orphanet J Rare Dis 13, 165.

Gandin, V., Brina, D., Marchisio, P.C., Biffo, S., 2010. JNK inhibition arrests cotranslational degradation. Biochim Biophys Acta 1803, 826–831.

Greer, P.L., Hanayama, R., Bloodgood, B.L., Mardinly, A.R., Lipton, D.M., Flavell, S.W., Kim, T.K., Griffith, E.C., Waldon, Z., Maehr, R., Ploegh, H.L., Chowdhury, S., Worley, P.F., Steen, J., Greenberg, M.E., 2010. The Angelman Syndrome protein Ube3A regulates synapse development by ubiquitinating arc. Cell 140, 704–716.

Inada, T., 2017. The Ribosome as a Platform for mRNA and Nascent Polypeptide Quality Control. Trends Biochem Sci 42, 5–15.

Kozlov, G., Denisov, A.Y., Girard, M., Dicaire, M.J., Hamlin, J., McPherson, P.S., Brais, B., Gehring, K., 2011. Structural basis of defects in the sacsin HEPN domain responsible for autosomal recessive spastic ataxia of Charlevoix-Saguenay (ARSACS). J Biol Chem 286, 20407–20412.

Krygier, M., Konkel, A., Schinwelski, M., Rydzanicz, M., Walczak, A., Sildatke-Bauer, M., Ploski, R., Slawek, J., 2017. Autosomal recessive spastic ataxia of Charlevoix-Saguenay (ARSACS) - A Polish family with novel SACS mutations. Neurol Neurochir Pol 51, 481–485.

Lariviere, R., Gaudet, R., Gentil, B.J., Girard, M., Conte, T.C., Minotti, S., Leclerc-Desaulniers, K., Gehring, K., McKinney, R.A., Shoubridge, E.A., McPherson, P.S., Durham, H.D., Brais, B., 2015. Sacs knockout mice present pathophysiological defects underlying autosomal recessive spastic ataxia of Charlevoix-Saguenay. Hum Mol Genet 24, 727–739.

Lariviere, R., Sgarioto, N., Marquez, B.T., Gaudet, R., Choquet, K., McKinney, R.A., Watt, A.J., Brais, B., 2019. Sacs R272C missense homozygous mice develop an ataxia phenotype. Mol Brain 12, 19.

Liutkute, M., Samatova, E., Rodnina, M.V., 2020. Cotranslational Folding of Proteins on the Ribosome. Biomolecules 10.

Longo, F., Benedetti, S., Zambon, A.A., Sora, M.G.N., Di Resta, C., De Ritis, D., Quattrini, A., Maltecca, F., Ferrari, M., Previtali, S.C., 2020. Impaired turnover of hyperfused mitochondria in severe axonal neuropathy due to a novel DRP1 mutation. Hum Mol Genet 29, 177–188.

Masciullo, M., Modoni, A., Tessa, A., Santorelli, F.M., Rizzo, V., D’Amico, G., Laschena, F., Tartaglione, T., Silvestri, G., 2012. Novel SACS mutations in two unrelated Italian patients with spastic ataxia: clinico-diagnostic characterization and results of serial brain MRI studies. Eur J Neurol 19, e77–78.

Michel, A.M., Fox, G., A, M.K., De Bo, C., O’Connor, P.B., Heaphy, S.M., Mullan, J.P., Donohue, C.A., Higgins, D.G., Baranov, P.V., 2014. GWIPS-viz: development of a ribo-seq genome browser. Nucleic Acids Res 42, D859–864.

Morimoto, R.I., Cuervo, A.M., 2014. Proteostasis and the aging proteome in health and disease. J Gerontol A Biol Sci Med Sci 69 Suppl 1, S33–38.

Muller, J.P., Scholl, S., Kunick, C., Klempnauer, K.H., 2021. Expression of protein kinase HIPK2 is subject to a quality control mechanism that acts during translation and requires its kinase activity to prevent degradation of nascent HIPK2. Biochim Biophys Acta Mol Cell Res 1868, 118894.

Parfitt, D.A., Michael, G.J., Vermeulen, E.G., Prodromou, N.V., Webb, T.R., Gallo, J.M., Cheetham, M.E., Nicoll, W.S., Blatch, G.L., Chapple, J.P., 2009. The ataxia protein sacsin is a functional co-chaperone that protects against polyglutamine-expanded ataxin-1. Hum Mol Genet 18, 1556–1565.

Pechmann, S., Willmund, F., Frydman, J., 2013. The ribosome as a hub for protein quality control. Mol Cell 49, 411–421.

Ricca, I., Morani, F., Bacci, G.M., Nesti, C., Caputo, R., Tessa, A., Santorelli, F.M., 2019. Clinical and molecular studies in two new cases of ARSACS. Neurogenetics 20, 45–49.

Romano, A., Tessa, A., Barca, A., Fattori, F., de Leva, M.F., Terracciano, A., Storelli, C., Santorelli, F.M., Verri, T., 2013. Comparative analysis and functional mapping of SACS mutations reveal novel insights into sacsin repeated architecture. Hum Mutat 34, 525–537.

Sato, S., Ward, C.L., Kopito, R.R., 1998. Cotranslational ubiquitination of cystic fibrosis transmembrane conductance regulator in vitro. J Biol Chem 273, 7189–7192.

Synofzik, M., Soehn, A.S., Gburek-Augustat, J., Schicks, J., Karle, K.N., Schule, R., Haack, T.B., Schoning, M., Biskup, S., Rudnik-Schoneborn, S., Senderek, J., Hoffmann, K.T., MacLeod, P., Schwarz, J., Bender, B., Kruger, S., Kreuz, F., Bauer, P., Schols, L., 2013. Autosomal recessive spastic ataxia of Charlevoix Saguenay (ARSACS): expanding the genetic, clinical and imaging spectrum. Orphanet J Rare Dis 8, 41.

Terracciano, A., Casali, C., Grieco, G.S., Orteschi, D., Di Giandomenico, S., Seminara, L., Di Fabio, R., Carrozzo, R., Simonati, A., Stevanin, G., Zollino, M., Santorelli, F.M., 2009. An inherited large-scale rearrangement in SACS associated with spastic ataxia and hearing loss. Neurogenetics 10, 151–155.

Thiffault, I., Dicaire, M.J., Tetreault, M., Huang, K.N., Demers-Lamarche, J., Bernard, G., Duquette, A., Lariviere, R., Gehring, K., Montpetit, A., McPherson, P.S., Richter, A., Montermini, L., Mercier, J., Mitchell, G.A., Dupre, N., Prevost, C., Bouchard, J.P., Mathieu, J., Brais, B., 2013. Diversity of ARSACS mutations in French-Canadians. Can J Neurol Sci 40, 61–66.

Vermeer, S., van de Warrenburg, B.P., Kamsteeg, E.J., Brais, B., Synofzik, M., 1993 updated 2020. Arsacs, in: Adam, M.P., Ardinger, H.H., Pagon, R.A., Wallace, S.E., Bean, L.J.H., Stephens, K., Amemiya, A. (Eds.), GeneReviews((R)), Seattle (WA).

Wang, F., Canadeo, L.A., Huibregtse, J.M., 2015. Ubiquitination of newly synthesized proteins at the ribosome. Biochimie 114, 127–133.

Wang, F., Durfee, L.A., Huibregtse, J.M., 2013. A cotranslational ubiquitination pathway for quality control of misfolded proteins. Mol Cell 50, 368–378.

Waudby, C.A., Dobson, C.M., Christodoulou, J., 2019. Nature and Regulation of Protein Folding on the Ribosome. Trends Biochem Sci 44, 914–926.

Willmund, F., del Alamo, M., Pechmann, S., Chen, T., Albanese, V., Dammer, E.B., Peng, J., Frydman, J., 2013. The cotranslational function of ribosome-associated Hsp70 in eukaryotic protein homeostasis. Cell 152, 196–209.

Wolff, S., Weissman, J.S., Dillin, A., 2014. Differential scales of protein quality control. Cell 157, 52–64.

Xiromerisiou, G., Dadouli, K., Marogianni, C., Provatas, A., Ntellas, P., Rikos, D., Stathis, P., Georgouli, D., Loules, G., Zamanakou, M., Hadjigeorgiou, G.M., 2020. A novel homozygous SACS mutation identified by whole exome sequencing-genotype phenotype correlations of all published cases. J Mol Neurosci 70, 131–141.

Zhou, M., Fisher, E.A., Ginsberg, H.N., 1998. Regulated Co-translational ubiquitination of apolipoprotein B100. A new paradigm for proteasomal degradation of a secretory protein. J Biol Chem 273, 24649–24653.

